# Direct Continuous EMG control of a Powered Prosthetic Ankle for Improved Postural Control after Guided Physical Training: a Case Study

**DOI:** 10.1101/2020.09.11.293373

**Authors:** Aaron Fleming, Stephanie Huang, Elizabeth Buxton, Frank Hodges, He (Helen) Huang

**Affiliations:** Joint Department of Biomedical Engineering, North Carolina State University and University of North Carolina at Chapel Hill, Raleigh, NC 27606, USA; UNC Hospitals, Department of Rehabilitation Therapies, Chapel Hill, NC 27514, USA; Prosthetic and Orthotic Fabrication, SunStone Lab LLC, Raleigh, NC 27615, USA

**Keywords:** Transtibial amputee, Postural Control, Myoelectric Control, Training, Dynamic Standing Balance

## Abstract

**Background:** Despite the promise of powered lower limb prostheses, the existing control of these modern devices is insufficient to assist many daily activities, such as postural control while lifting weight, that require continuous control of prosthetic joints according to human states and environments. The objective of this case study was to investigate the feasibility and potential of direct, continuous electromyographic (dEMG) control of a powered ankle prosthesis, combined with physical therapist (PT)-guided training, for improved standing postural control in an individual with transtibial amputation.

**Methods:** A powered prosthetic ankle was directly controlled by EMG signals of the residual *lateral gastrocnemius* and *tibialis anterior* muscles. The participant with transtibial amputation received 4-week PT-guided training on posture while using the dEMG control of powered ankle. A subset of activities in the mini-BESTest (a clinical balance assessment tool) were used in the training and evaluation protocol. We quantified EMG signals in the bilateral shank muscles, biomechanics that captures postural control and stability, and score for the clinical balance evaluation.

**Results:** Compared to the participant’s daily passive prosthesis, the dEMG-controlled ankle, combined with the training, yielded improved clinical balance score and reduced compensation from the intact joints. In addition, cross correlation coefficient of bilateral CoP excursions, a metric for quantifying standing postural control, increased to *0.83(±0.07)* when using dEMG ankle control, compared with *0.39(±0.29)* when using the passive device. Between-limb coordination was also observed as synchronized activation of homologous muscles in the shank. We witnessed rapid improvement in performance on the first day of the training for load transfer tasks, where bilateral CoP synchronization improvement was significantly related to repetition order (*R=0.459, p = 0.045)*. Finally, the participant further improved this performance significantly across training days.

**Conclusion:** This case study showed the feasibility of dEMG control of powered prosthetic ankle by a transtibial amputee after a PT-guided training to assist postural control. This study’s training protocol and dEMG control method that lays the foundation for future study to extend these results through the inclusion of more participants and activities.

## Introduction

Recent advances in intelligent, powered prosthetic legs have opened up opportunities for individuals with lower limb amputations to restore their normative walking patterns on uneven terrains [1–9]. These modern devices use primarily autonomous control, which is, however, inadequate to assist other important daily tasks that involve unpredictable, non-cyclic motor behavior and require continuous coordination with the user’s motor control and environments. One example of such activities is anticipatory and compensatory postural control in standing, walking, or other recreation activities [10, 11].

Focusing on standing postural control, lower limb amputees wearing passive prostheses have shown decreased postural stability and increased compensation from the intact limb [12, 13]. This is partly because of the lack of active degrees of freedom in the prostheses. Powered prostheses have active, controllable joints and, therefore, a potential to enhance the amputees’ postural stability. Unfortunately, there has been no autonomous control solutions to assist amputees’ standing posture because it is difficult to predict the postural perturbations and human motor control strategy for counteracting the perturbations. We are aware of only one research group, developing autonomous prosthesis control to assist posture stability of the prosthesis users when standing on slops [14]. The controller detected the inclination angle after the prosthesis foot was on a slope and then adjusted equilibrium position of prosthesis joint in order to support the amputee’s standing posture. This automated control was reactive and limited in function because it can assist standing posture on a slope only, and it acted only after the prosthesis foot was on an incline. Hence, this prosthesis control was insufficient to assist anticipatory postural control (i.e., action before the perturbation happens) or handle the postural control under dynamic perturbations (e.g. weight transfer), which requires continuous postural control based on the shift of center of mass (COM).

As human neural control system is highly adaptable to the task context, perhaps neural control of prosthetic joint can be a viable solution to assist the amputee’s postural control and balance stability. EMG signals of the residual muscles are readily-available efferent neural sources in amputees and has been used for neural control of prosthetic legs in walking. Many groups have used EMG pattern recognition to classify the user’s locomotor tasks, switching autonomous prosthesis control mode accordingly for enabling seamless locomotor task transitions [15–19]. Another group used EMG signal magnitude recorded from the residual *medial gastrocnemius* (GAS) to proportionally modulate a control parameter in the automated prosthesis control in the push-off phase [20]. Both aforementioned approaches relied on autonomous prosthesis control laws and cannot produce neural control of prosthetic joints continuously. Three other groups conducted case studies to show the feasibility of direct EMG (dEMG) control in walking, in which EMG magnitude of one or a pair of residual antagonistic muscles are directly mapped to modulate the applied torque to the prosthetic joints continuously [21–23]. In dEMG control, the behavior of prosthetic joints is primarily and continuously determined by human neuromuscular control, mimicking human biological joint control. Note that the existing studies on EMG control of powered prosthetic legs, regardless the methods used, focuses on locomotor tasks mainly. Little effort has been aimed to address postural control.

One of the main challenges in implementing direct EMG control of a prosthesis, although technically simple, is whether amputees are capable of producing coordinated activations between the residual antagonist muscles for operating a prosthesis joint in dynamic task performance.

Previous studies have shown a large variation among transtibial amputees in producing coordinated activity between the residual *tibialis anterior* (TA) and GAS in a sitting posture or walking [22, 24, 25]. These results implied that individuals with transtibial amputations might no longer manifest normative activation in the residual muscles due to the limb amputation. Luckily, evidences have also shown that training or practice is a potential way to improve the capability of amputees in modulating residual muscles’ activity for dEMG control. Our previous study [26] tested transtibial amputees in dEMG control of a virtual inverted pendulum, mimicking the dynamics of standing posture. We noted improved task performance for all the amputee participants after a short-term practice within the same experimental visit. However, the amount of improvement varied significantly among the participants. Acclimation to dEMG control has involved repeating the evaluated task (walking) for an extend period of time [21, 27], or visualizing phantom limb movements [28]. For Huang et al. [27] transtibial amputees did not adapt activation of their residual GAS until they were given visual feedback of their prosthetic ankle angle with a target trajectory. However, it was unclear whether, after removing biofeedback training, amputees could still reproduce desired ankle joint trajectories or continue to improve control. Dawley et al. [21] observed improvements residual muscle activity after simply repeated walking sessions with a single above-knee amputee. From the findings of previous studies we postulate that amputees might adapt and learn the necessary muscle activation pattern for control function after training and practice. We expand the work of previous studies by 1) Creating and implementing an four-week PT-guided training paradigm without supplementary feedback of the ankle prosthesis 2) Implementing activity from *both* TA and GAS residual muscles for direct, continuous prosthetic ankle control and 3) Investigating the ability for an amputee to improve standing postural control with this control paradigm.

In this paper we present a case study to demonstrate the feasibility and potential benefit of dEMG control of a powered ankle prosthesis on an individual with a transtibial amputation for enhanced postural stability. Since training is likely necessary for successful application of dEMG control, the case study included four-week of physical therapist (PT)-guided training. Via this case study, we are interested in learning (1) how a transtibial amputee learns residual muscle activation patterns and coordination necessary for driving a powered ankle prosthesis for postural control, and (2) whether dEMG control of a powered ankle can improve the postural stability of transtibial amputees, compared to the currently prescribed passive prostheses. The results may inform the needed training protocol for dEMG control of prosthetic ankle and the future development of versatile powered prostheses that can assist various activities of individuals with transtibial amputations.

## Materials and Methods

### Participant

We recruited one amputee participant to take part in this case study. The participant provided informed, written consent to participate in this Institutional Review Board approved study at the University of North Carolina at Chapel Hill. The participant was 57 years old and 3 years post-amputation with septic shock as the cause. The participant weighed 131kg. The participant used a pin-lock suspension and a Pro-Flex foot (Össur) daily. For the purpose of the study the participant was fit with a new prosthetic socket (StabileFlex, Coyote Design). This transtibial socket design provided more room in the anterior-posterior direction while still maintaining adequate fit by loading the medio-lateral sides of the residual limb more heavily. This socket design provided more room for the residual *tibialis anterior* and residual *gastrocnemius* muscles to contract compared to traditional socket designs, which increased comfort of residual muscle contractions within the socket and reduced residual muscle fatigue. On a daily basis the participant used his passive prosthesis for household and community ambulation. He was able to traverse environmental barriers without requiring an assistive device and was independent with daily tasks, including driving.

As a case study we sought to understand how an amputee with relatively average control of his residual muscles would perform in this extended training with a dEMG controlled prosthesis. Thus, we recruited this participant based on his EMG control performance with his residual antagonistic muscles in a previous study [26] (participant TT2).

### Clinical Screening

We conducted a sensory screening of the participant before the start of the study. A trained physical therapist performed a sensation screen of the participant’s residual and intact limb. We noted partial neuropathy in the participants intact foot. The participant had diminished light touch sensation distal to the ankle joint. The participant had absent light touch sensation at the medical aspect of the intact foot. Above the ankle joint, the participant was able to localize light touch sensation stimuli in all dermatomes bilaterally.

### Device and Setup

We used an experimental ankle prosthesis driven by pneumatic artificial muscles (PAM) as the platform for testing proportional myoelectric control (dEMG control) with residual muscles. Technical information about the device can be found [29]. This device was initially developed for the study of continuous, proportional myoelectric control of plantarflexion during level-ground walking [27]. In this study we implemented continuous control of both dorsi- and plantar-flexion using two sets of proportional pressure valves (MAC Valves, Wixom, MI, USA) with two valves allocated to each PAM for a total of 8 valves. The input control signal for the control valves was 0-10V which corresponded to a pressure output of 0-90psi proportionally.

We processed electromyographic (EMG) signals from residual *tibialis anterior* (TA) and residual *lateral gastrocnemius* (GAS) muscle in real-time (dSPACE, CLP-1103, 0-10V output) to create a smoothed control signal for each set of pressure valves. The real-time setup created a smoothed control signal by first applying a high-pass filter (100Hz, 2nd Order Butterworth) to reduce the effect of potential signal artifacts. The setup then rectified the signal and applied a low-pass filter (2Hz, 2nd Order Butterworth). The smoothed signal for each respective muscle was then sent to the pressure regulators, which generated pressure proportionally to the input voltage to actuate the device.

We applied a baseline signal from the setup for both pairs of muscles to set a base stiffness for the ankle prosthesis. While the dEMG control was off, and the prosthesis unloaded, we applied a baseline signal that generated a neutral ankle position (5-7 degrees dorsiflexion). We then asked the participant to stand and we adjusted baseline signals based on the perceived baseline stiffness compared with his intact ankle. After iterating this process, we established a baseline signal of ~3V for the plantar- and dorsi-flexor muscles. When the participant had active control (dEMG control was turned on) we observed an average tonic activity from the residual muscles (~1.3V from residual GAS, ~1V from residual TA) across sessions. In order to allow the participant true continuous control of the prosthetic device we did not enforce an EMG threshold that would restrict low-level activity from controlling the device. We applied a gain to each control signal at the beginning of each session in order that a maximum contraction generated a control signal between 9-10V.

### Introduction to the system

Before the initial evaluation and training, we introduced the amputee participant to the direct EMG control paradigm and the pneumatic ankle device. While sitting, the participant wore the powered ankle prosthesis and was given time to freely move the ankle joint via residual muscle contractions. During this free exploration we provided visual feedback of his residual muscle activation as a percentage of his maximum voluntary contraction (%MVC). In order to facilitate learning the dynamics (i.e. possible combinations of ankle joint stiffness) we then asked the participant to fill a virtual control input space with his residual antagonistic muscle contractions (as described in [30]). We then repeated these steps while the amputee participant stood with handlebar support available to him. We took these steps to provide the participant with a clear understanding of the input-output relationship of reciprocal activation and co-activation of his residual muscles to changes prosthetic ankle joint dynamics. After this introduction stage we did not provide the amputee participant visual feedback of residual muscle activations.

### Training and Evaluation Sessions

The study consisted of an initial evaluation, 5 training sessions, a final evaluation, and a supplementary evaluation. The timeline for training and evaluation sessions are outlined (Table 1).

For the evaluation sessions we asked the participant to perform quiet standing tasks across various sensory conditions. The four tasks selected involve quiet standing under two visual conditions, Eyes Open (EO) and Eyes Closed (EC), and two surface conditions, Firm and Foam, as described by the BESTest [31]. These tasks were scored by a trained physical therapist on a scale from 0-3 where when the participant stood stably for 30 seconds (score = 3), 30 seconds unstable (score = 2), stood less than 30 seconds (score = 1), and unable (score = 0) [31].

**Table 1.** Clinical Standing Balance Evaluation and Training Timeline

| Day | Day 1 | Day 2 | Day 3 | Day 4 | Day 5 | Day 6 | Day 7 | Day 8 |
| --- | --- | --- | --- | --- | --- | --- | --- | --- |
| Session Type | Passive Prosthesis & dEMG Prosthesis Evaluation* | Training (dEMG only) |  |  |  |  | dEMG Evaluation | Supplementary Eval. (Passive only) |
| Tasks | Quiet Standing:<br>1) Firm, EO**<br>2) Firm, EC<br>3) Foam, EO<br>4) Foam, EC | Rocker Board Warm-up<br>Arm Raise<br>Forward Reach<br>Load Transfer<br>Sit-to-Stand |  |  |  |  | Quiet Standing:<br>1) Firm, EO<br>2) Firm, EC<br>3) Foam, EO<br>4) Foam, EC | Rocker Board Warm-up<br>Arm Raise<br>Forward Reach<br>Load Transfer<br>Sit-to-Stand |

\* Passive prosthesis evaluation conducted first \*\* EO: Eyes Open, EC: Eyes Closed

For the training sessions we selected tasks relevant to daily life activities: Load transfer, Sit-to-Stand, Forward reach, and Arm raise. These tasks (with the exception of the load transfer) are also a subset of evaluation tasks in the BESTest [31]. We selected these training tasks to differ from the evaluation tasks in order to understand the effect of training to overall standing stability, as opposed to task-specific stability, while using the dEMG control of a prosthetic ankle. At the start of each training session we asked the amputee to stand with his prosthetic foot on a rocker-board and intact foot on firm ground for 30 seconds. During training the participant completed 2 trials of each task per session, with a minimum of 4 repetitions per trial. The number of repetitions increased across days, as prescribed by the physical therapist, where day 4 of the training sessions (Table 1) had 20 total repetitions of each task.

We conducted the study over the course of 25 days. We gave a minimum of 1 day of rest between sessions in order to reduce fatigue effects and a maximum of 4 days of rest between sessions to minimize learning losses. We conducted training with the dEMG controlled device only. We evaluated standing stability with both passive and dEMG controlled devices on the first day. After training, we performed a follow-up evaluation with the dEMG control. In order to compare postural control strategies in training tasks across devices we conducted a supplementary evaluation session where the participant repeated the training tasks while wearing his passive device.

A trained clinician attended each training session with the participant. During each training session the clinician observed the participant complete each task. Between repetitions, the clinician provided feedback to the participant regarding his full-body symmetry, body mechanics, foot positioning, and alignment. The clinician provided feedback to encourage equal contribution from both limbs toward the specific task. The patient received verbal cues to shift his weight onto his prosthetic side and to recruit muscles in a “toes up” or “toes down” direction when learning each task. This directional cue is the same language used when he performed his warm-up on the rocker board. He also required cues to shift his weight onto his prosthetic side, especially for tasks such as sit to stand transfers in which he was accustomed to compensating for an ankle that was relatively fixed, whereas the power prosthesis allowed for movement in the sagittal plane.

### Measurements

During all sessions we recorded activity from the residual and intact shank muscles. Specifically, we placed EMG sensors (Neuroline 715, 1mm height) on residual *lateral gastrocnemius* and residual *tibialis anterior* muscles (Figure 1). We located residual muscle bellies via palpation while the participant contracted and relaxed his muscles [32]. We then routed cables away from bony landmarks and connected them to a pre-amplifier (Motion Lab Systems, MA-412, Gainx20) outside of the prosthetic socket. We placed EMG sensors (Motion Lab Systems, MA-420, Gainx20) on intact GAS and intact TA muscles. We connected all sensors to an amplifier (MA300-XVI, Gain x1000).

**Figure 1.**
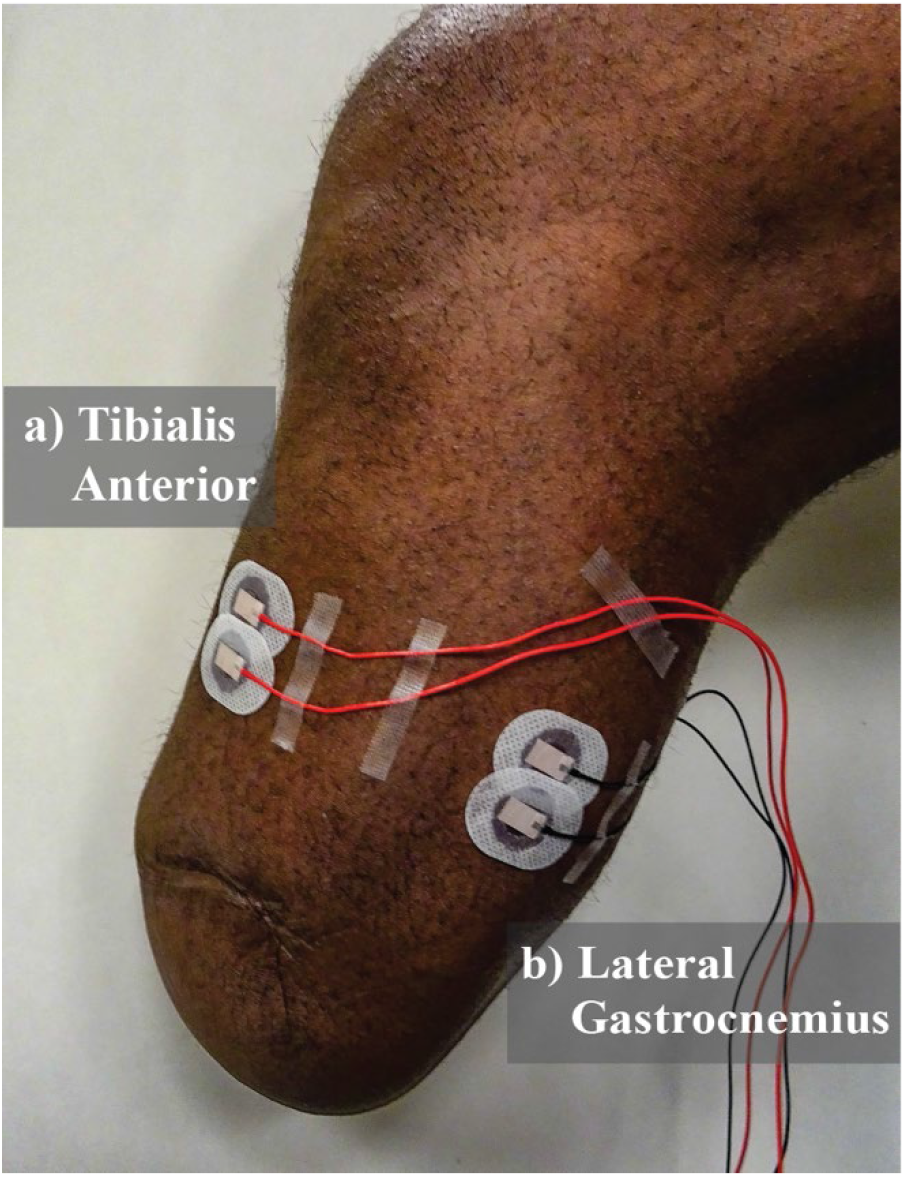
Residual limb electrode placement. a) Tibialis Anterior electrodes. b) Lateral Gastrocnemius electrodes. Electrodes are placed in line with muscle belly (location determined through palpation as amputee is asked to contract muscle). Cables are routed away from bony landmarks.

For all sessions we collected Center of Pressure (CoP) locations under each foot using an instrumented split-belt treadmill (1000Hz, Bertec Corp.). For the final session of training (Day 6) and the supplementary passive evaluation session (Day 8) we collected full-body kinematics using motion capture (100Hz, 53 markers, VICON, Oxford, UK).

### Data Analysis

We processed all data offline using Matlab (Mathworks, Natick, MA). We analyzed all quiet standing trials where the participant was able to maintain balance for the entire trial without stepping. Since the participant was unable to maintain balance in the dEMG control, Pre-Training, Foam condition, we used the score given by the physical therapist for comparison. For the training sessions and supplementary evaluation, we analyzed data from the load transfer tasks only. We selected the load transfer task for analysis since this was self-reportedly the most difficult task for the participant during training.

For the training session analysis, we extracted and evaluated each repetition of the load transfer task. Each repetition was manually extracted through visual inspection of the summed vertical ground reaction forces in order to determine the moment the weight was picked up (before pick-up the weight was located beside the instrumented treadmill). Based on the speed of movement during training we empirically windowed each repetition to ±2s on either side of the moment of pick-up.

For all evaluation trials and load transfer repetitions we calculated synchronization of CoP excursions in the Anterior-Posterior direction under each foot by taking the cross-correlation between the time series [33]. For each trial, we subtracted the mean CoP values from each foot and conducted a cross-correlation of the times series using MATLAB (xcorr). We determined the cross-correlation coefficient at time zero (CC_0_), max cross-correlation coefficient (CC_max_), and the lag value (Lag_CC_) in milliseconds from time zero to CC_max_. CC_max_ and Lag_CC_ are calculated to determine potential lag in CoP excursions between limbs using a window of ±1s [33].

For the final training session (dEMG control) and in the supplementary session (passive) we analyzed ankle, knee, and hip joint flexion during the load transfer task. We calculated joint angles in the sagittal plane [34] for each windowed repetition. We then subtracted joint angles during quiet standing from all repetitions for each condition. We tabulated peak hip, knee, and ankle flexion angles during the windowed repetitions.

In order to analyze the neural control strategy used by the participant we processed EMG activity from residual and intact TA and GAS muscles. We first high-pass filtered the raw EMG signal (Butterworth, 2^nd^ order, 100Hz cutoff) from all muscles to remove potential motion artifacts. We rectified the signals and applied a low-pass filter (Butterworth, 2^th^ order, 20Hz cutoff) in order to generate a smoothed signal for qualitative comparison. We then selected representative repetitions from the first and final day of training based on CC_0_ values that were closest to the average CC_0_ for that day of training. We then plotted CoP excursion, EMG activity from residual and intact TA and GAS, and residual TA and GAS control signals together for qualitative comparison.

### Statistical Analysis

For our statistical analysis of the data we used the statistical software (JMP, SAS, US). We used a one-way ANOVA to compare the CC_0_, CC_max_, and Lag_CC_ with Training Day as the main effect. We used the Shapiro-Wilk normality test (p<0.01) to detect outlier repetitions. One repetition was removed from our analysis (repetition 2, day 1 training, CC_0_ = −0.4). We ran a simple linear regression to determine the amount of variance (via R-squared) described by trial order in each training session CC_0_, CC_max_, and Lag_CC_. We analyzed joint flexion angles in the load transfer task between dEMG control and passive device. We used a two-way ANOVA to detect main and interaction effects of Device and Joint. When we found a significant effect, we tested for statistical differences within joint and device conditions using Tukey’s honestly significant difference test (*a* = 0.05). Significance thresholds were set using an alpha value of 0.05.

## Results

### Quiet Standing Evaluation: Clinical Scoring of Stability

We observed clear improvements in stability with the dEMG control of the powered ankle in the quiet standing tasks post-training (Table 2). In the pre-training condition, the amputee displayed moderate instability on the firm surface for both eyes open and eyes closed, evidenced by visually noticeable sways (*score = 2*). In the foam surface the amputee was unable to maintain balance without stepping in either condition (*score = 1*). Post-Training, the amputee improved stability over all conditions (*score = 3*). In all surface and vision conditions the participant did not display visually significant sways and did not require the use of any handlebars.

**Table 2:**
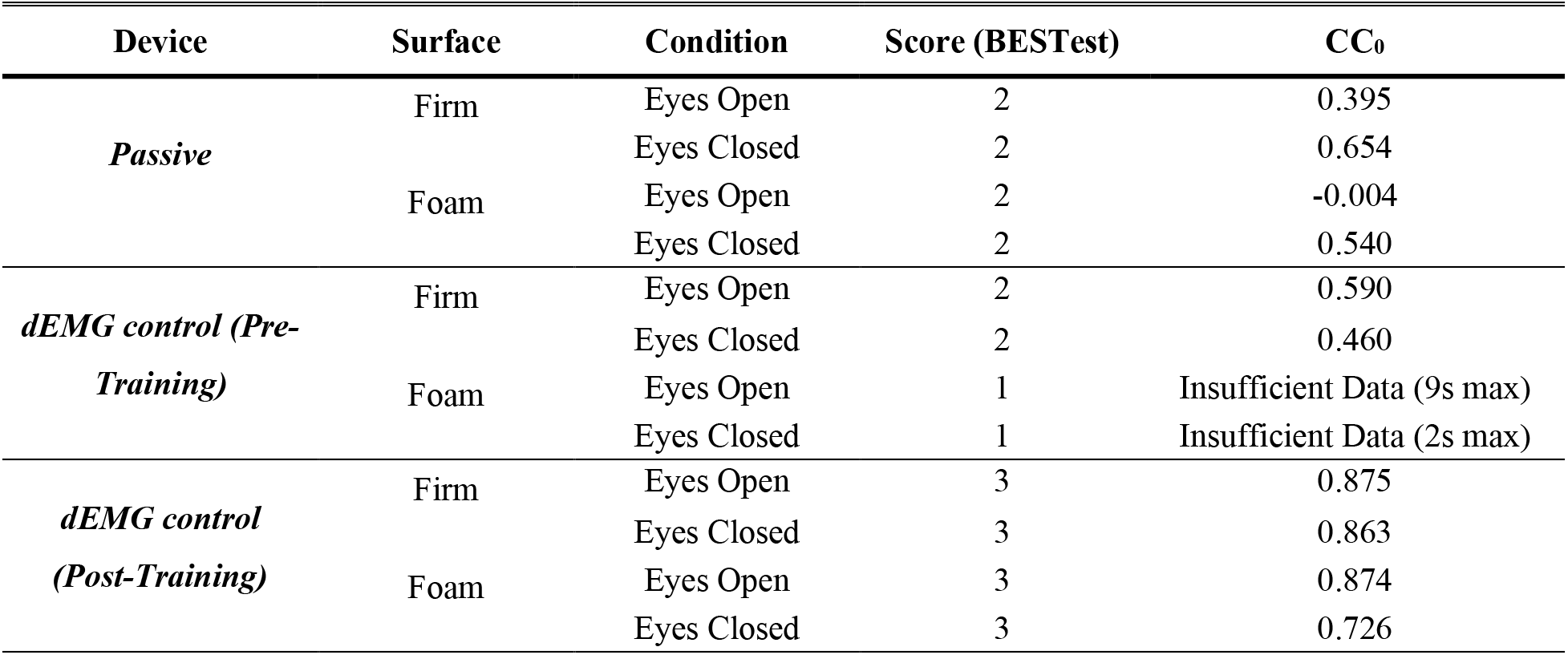
Quiet Standing Tasks Clinical Score and Between Limb Synchronization

We observed differences in stability between the passive (baseline) and dEMG controlled condition (Post-Training) (Table 2). With his passive device the amputee was able to maintain balance in all conditions with significant postural sway, and the use of handlebars was not needed (*score = 2*). With dEMG control, post-training, the amputee had minimal postural sways for all conditions (*score = 3*).

### Quiet Standing Evaluation: Between-Limb Synchronization

The participant demonstrated distinct patterns of bilateral center of pressure excursions between the passive and dEMG control (Post-Training) for the quiet standing tasks. Figure 2 shows this stark contrast in the foam condition where the participant displayed noticeably higher synchronization between his intact and prosthetic foot CoP_AP_ excursion with dEMG control (EO CC_0_ = 0.874, EC CC_0_ = 0.726) compared with his passive device (EO CC_0_ = 0.004, EC CC_0_ = 0.540). We observed this increase in synchronization during dEMG control in firm surface conditions as well (Table 1). The magnitude of CoP_AP_ excursion of the prosthetic foot in the passive device was less than the intact limb CoP_AP_ excursion as evidenced by time series plots (Figure 2a,b). The participant increased CoP_AP_ excursion on the prosthetic side post-training with dEMG control (Figure 2c,d).

**Figure 2:**
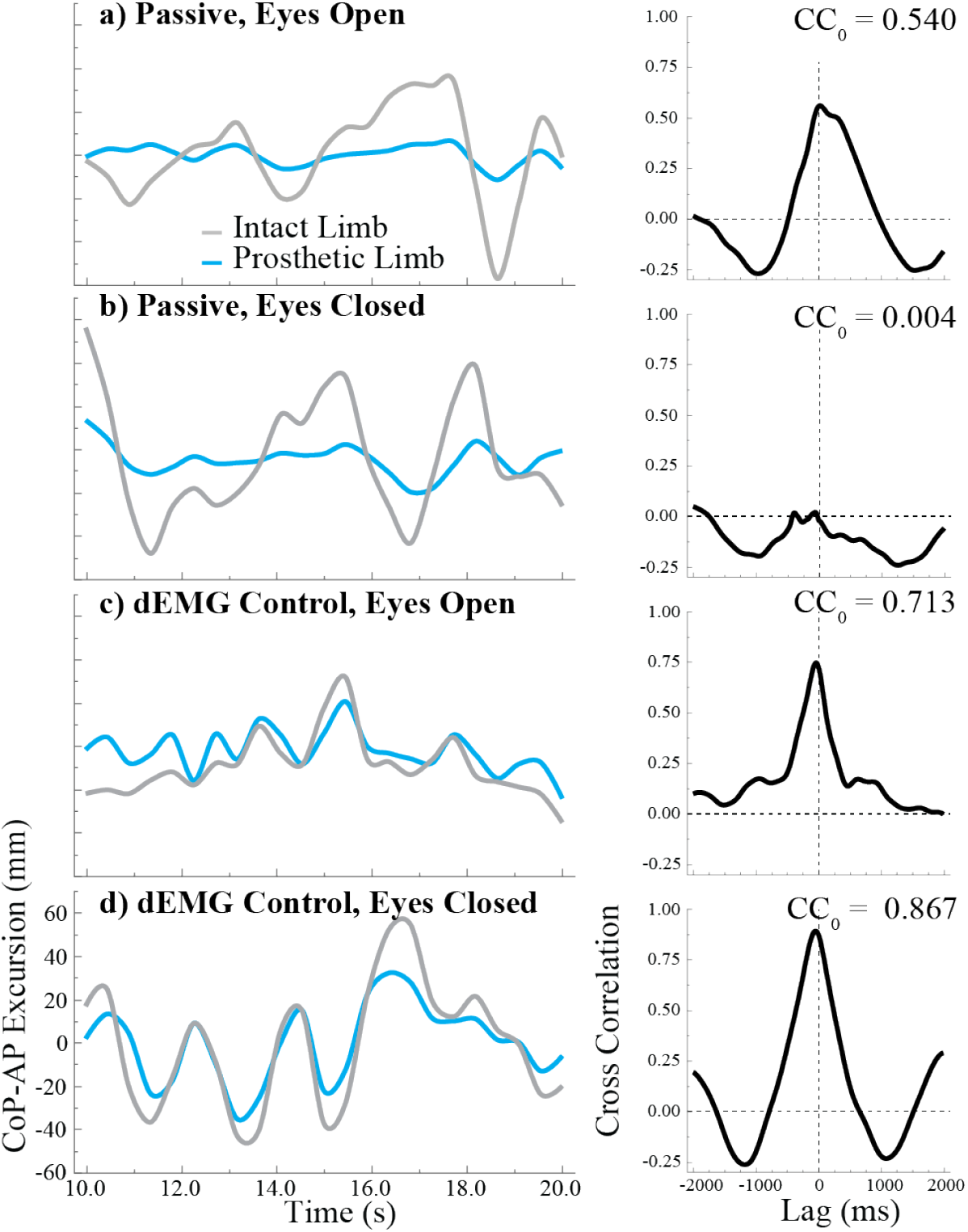
Passive vs. Post-training dEMG control on the Foam Surface. Representative center of pressure excursion and cross correlation between limbs. Representative trials are 10 second portions taken from each 30-second trial. Trials shown above are firm surface only. a) Passive device, eyes open condition b) Passive device, eyes closed condition c) dEMG controlled device, eyes open d) dEMG controlled device, eyes closed.

In dEMG control, the amputee demonstrated improvements in between limb synchronization after training for all quiet standing conditions (Table 2 & Figure 3). In the firm condition, pre-training, we observed moderate cross-correlation in CoP excursions between the intact and dEMG controlled foot (EO CC_0_ = 0.590, EC CC_0_ = 0.460) (Figure 3a,b). Post-training, the participant more closely synchronized CoP excursions between the two feet (EO CC_0_ = 0.875, EC CC_0_ = 0.863) (Figure 3c,d) in the firm condition. During the initial evaluation the amputee was unable to maintain balance in the foam condition thus we did evaluate CC_0_ for the pre-training, dEMG control condition. However, the amputee demonstrated similar synchronization values between firm and foam conditions in the post-training condition (*foam*: EO CC_0_ = 0.874, EC CC_0_ = 0.726).

**Figure 3:**
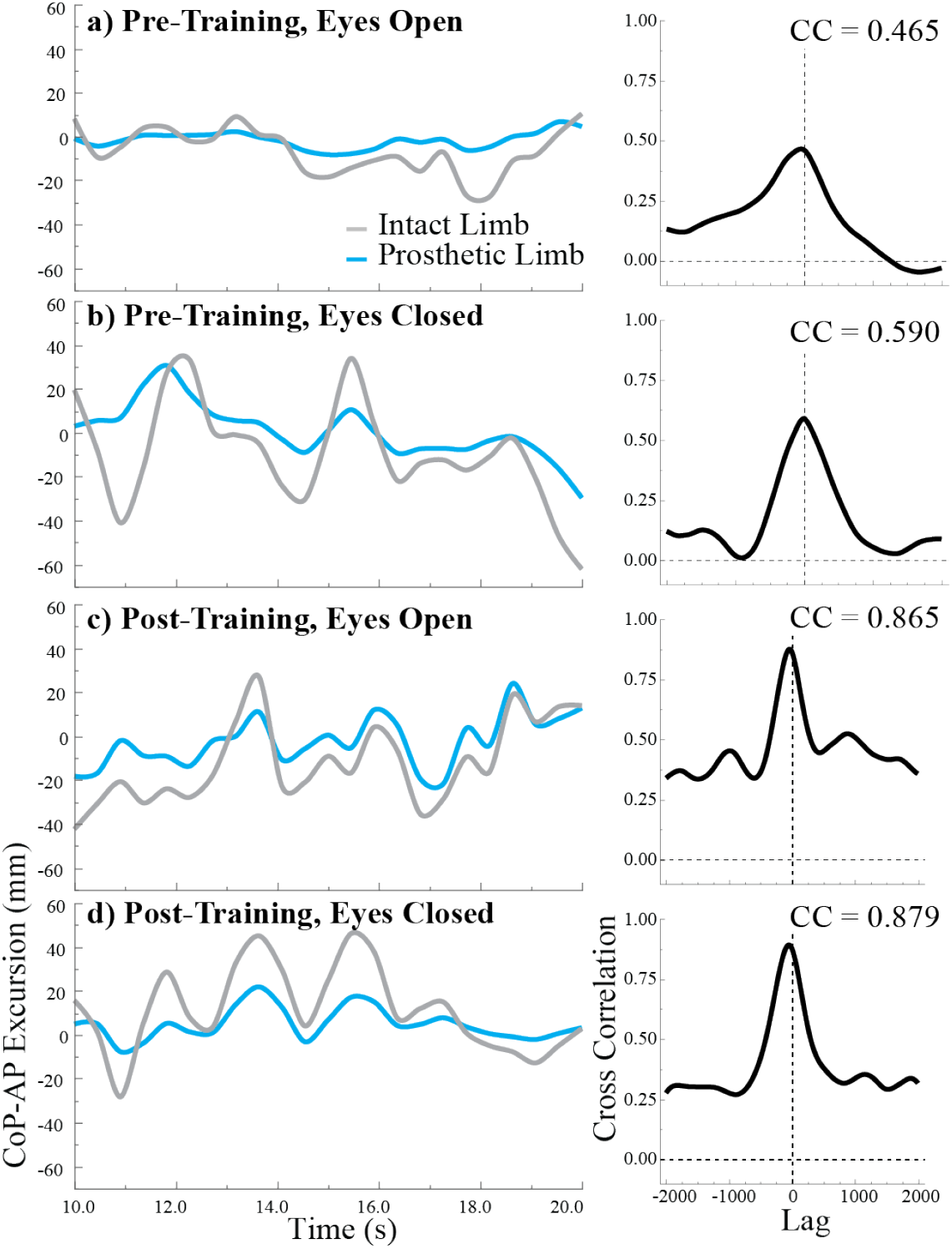
Pre vs. Post-training with dEMG control on the Firm Ground. Representative Center of Pressure excursion and Cross Correlation between limbs. Representative trials are 10 second portions taken from each 30-second trial. Trials shown above are firm surface only. a) Pre-training, eyes open condition b) Pre-training, eyes closed condition c) Post-training, eyes closed condition d) post-training, eyes closed condition.

### Training Evaluation: Load Transfer Task

Over the course of training the amputee significantly improved between-limb synchronization of CoP excursion. In the initial trials of the load transfer task, the participant displayed moderate levels of synchronization (CC_0_ = 0.49(±0.16), CC_max_ = 0.52(±0.14), CC_lag_ = −107.3ms (±357.1)) (Figure 4) similar to synchronization values observed during the initial evaluation. We observed that CC_max_ and CC_0_ improved significantly over the course of just the first day, where CC_max_ and CC_0_ are significantly related to repetition order (CC_max_: R^2^ = 0.459, p = 0.045; CC_0_: R^2^ = 0.646, p = 0.009) (Figure 4). We determined this relationship was significant for the first day, however not for the trial order in the remaining days. Across training, day was found to be a significant main effect for CC_max_ (*p = 0.011*) and CC_0_ (*p = 0.006*), but not for CC_lag_ (*p = 0.279*). At the final day of training we observed CC values of (CC_0_ = 0.76(±0.15), CC_max_ = 0.76(±0.16), CC_lag_ = −22.8ms (±32.79)).

**Figure 4.**
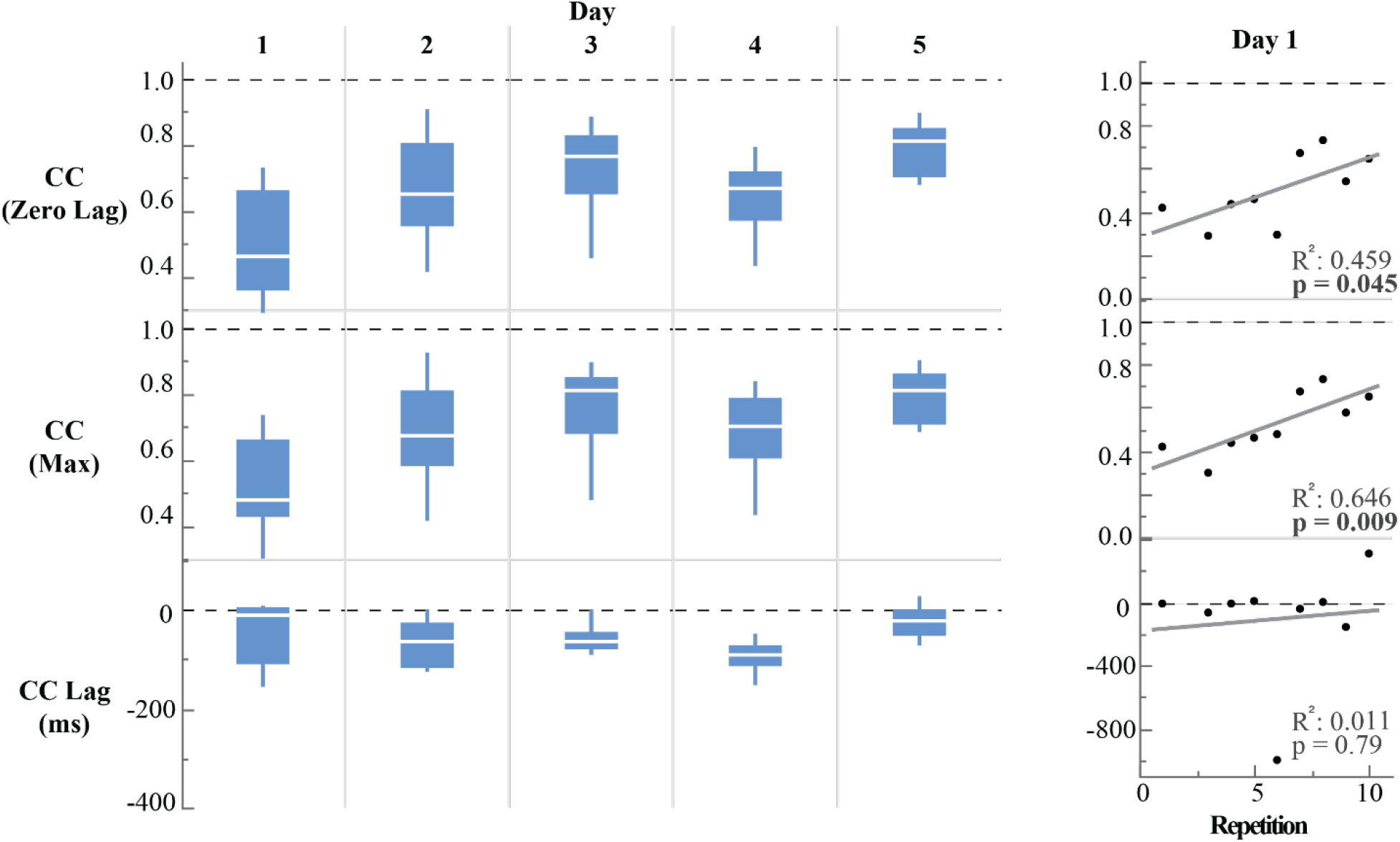
CoP synchronization values during training for the load transfer task. R-squared values and p-value are shown for Cross-Correlation (CC) values (CC at zero lag, maximum CC, and lag of maximum CC from zero lag) for day 1 of training. Due to concern for residual muscle fatigue during training, Day 1 and 2 contained less than 10 repetitions.

Analysis of EMG patterns during representative load transfers demonstrated distinct neural strategies between initial and final trials (Figure 5). These specific repetitions were chosen since their CC_0_ value closely matched average CC_0_ values for the initial and final day of training. Pre-training, we observed different timing and shape of EMG activity between the residual and intact limb. The participant intermittently activated the intact TA (Figure 5a) with a steady contraction of the GAS muscle throughout the movement (Figure 5b). In comparison the amputee had little to no activation from the residual TA before peak squat depth (Figure 5a,c) followed by significant activation of the GAS while returning to the standing posture (Figure 5b,c). The control signal reached half of its force generating potential (5V ~ 50psi) in the plantar-flexor direction during this movement (Figure 5c). Post-training, the strategy between the two limbs appeared more closely aligned. Activations from the residual TA were seemingly identical to activations from the intact TA (Figure 5e). Intact and residual GAS muscle activations were relatively aligned (Figure 5f) with the exception of activation of the intact GAS muscle before reversal of the squatting motion (Figure 5f). The control signal to the prosthesis (Figure 5c,g) mostly clearly demonstrated residual antagonistic pair control strategy across training. In final trials we observed high activations of the residual TA at the beginning of the movement, followed by small contractions from the residual GAS and co-contraction post-squat (Figure 5g). CC of CoP_AP_ excursions demonstrate the similarity in control strategy between limbs (Figure 5h).

**Figure 5:**
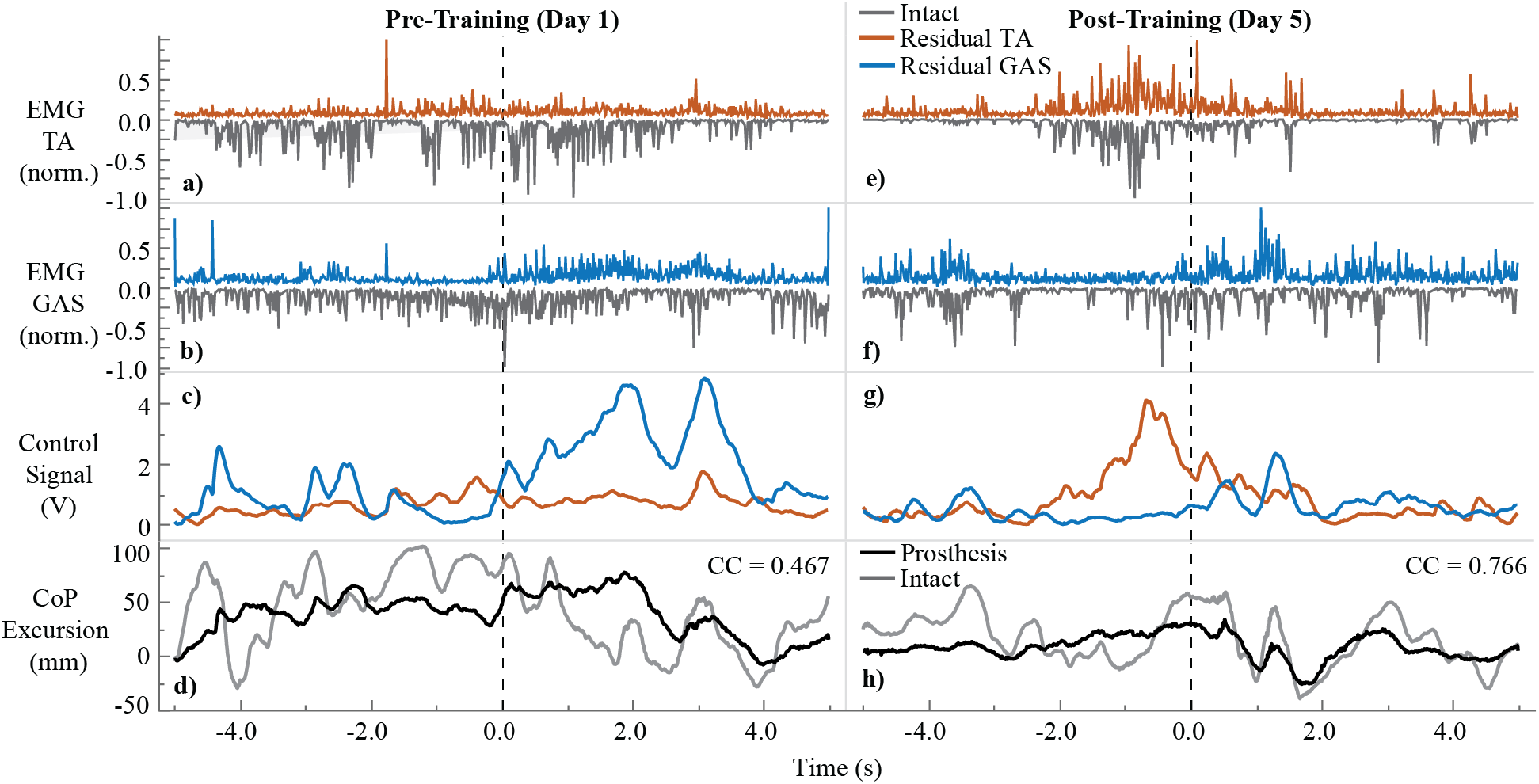
Representative load transfer trials pre and post-training. Dashed Line) moment of peak deceleration during squatting movement. **a)** normalized EMG of residual (orange) and intact (grey) TA muscle pair. **b)** Normalized EMG of residual (blue) and intact (grey) GAS muscle pair. **c)** Control signal to the prosthesis from the real-time processing of residual TA (orange) and residual GAS (blue) muscle EMG. **d)** CoP excursion from prosthetic (black) and intact foot (grey). Cross-correlation values are displayed for each representative trial (Pre: CC = 0.467, Post: CC = 0.766). **e-h)** Data for post-training. Normalized EMG was calculated by dividing the maximum EMG value for each muscle from the entire trial.

### Load Transfer Task: Postural Control Strategy

Post-training, we observed significantly different postural strategies between the passive and dEMG controlled device for the load transfer task. We observed small flexion angles for the passive ankle prosthesis during the load transfer (Table 3 & Figure 6). With dEMG control post-training, the ankle flexion angle significantly increased (Passive-dEMG, *p < 0.0001*). For the dEMG control condition the knee flexion angle also increased (Passive-dEMG, *p < 0.0001*) and the hip flexion angle decreased (Passive-dEMG, *p < 0.0001).* We observed a significant interaction between the device and joint (*p* < 0.0001).

**Table 3:**
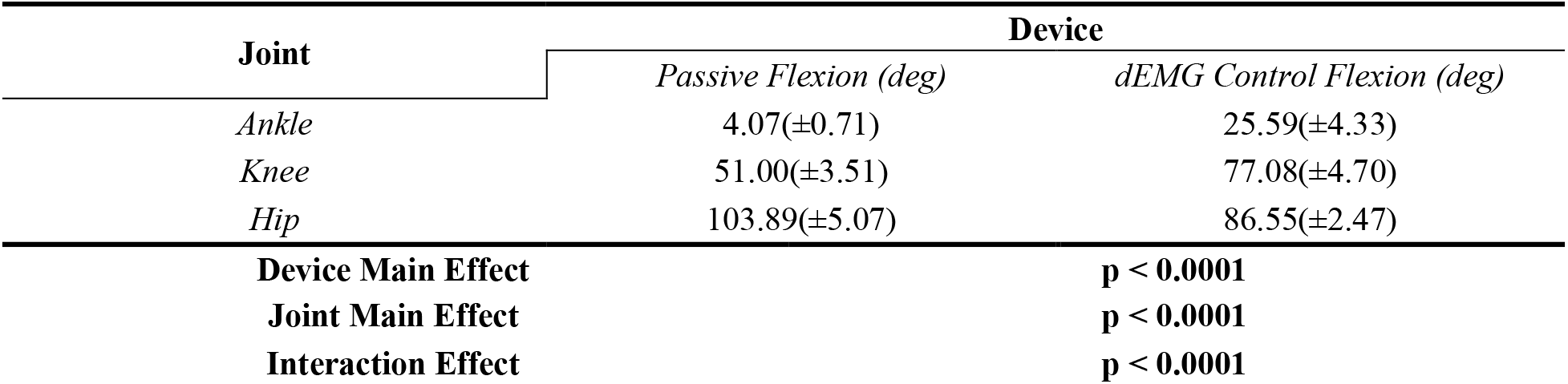
Load Transfer Joint Angle (Passive vs. Post-Training dEMG Control)

**Figure 6.**
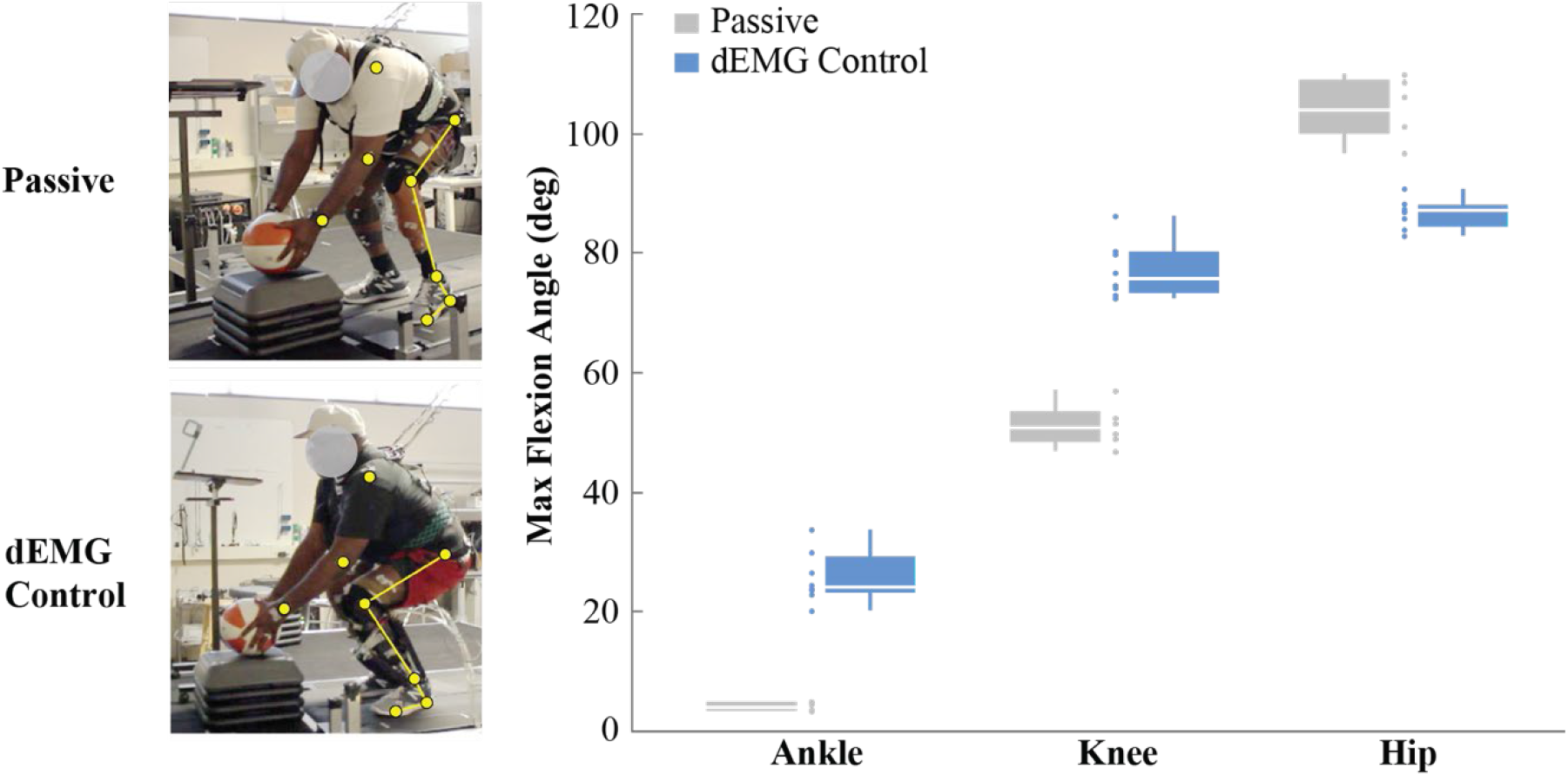
Load transfer task joint flexion angles (Passive vs. Post-Training dEMG Control). *Grey)* Passive prosthetic ankle, hip, and knee joint flexion on affected limb at peak squat depth (as determined by location of hip joint center). *Blue)* dEMG controlled prosthetic ankle, hip, and knee joint flexion at peak squat depth. Joint flexion angles determined as the difference between angle at maximum depth and angle during quiet standing.

## Discussion

In this study, we present the feasibility of direct EMG control to continuously operate prosthetic ankle joint mechanics in order to address the postural stability for individuals with transtibial amputations. The main finding of this study is that our recruited transtibial amputee participant was capable of using residual antagonist muscles to directly and continuously control a prosthetic ankle to significantly improve standing postural control, after 4 weeks of PT-guided training sessions, across various contexts compared with postural control using a passive ankle prosthesis as the baseline after. Completely different from the “standard” control framework for active lower limb prostheses and exoskeletons as suggested in [35] that relies on preprogrammed, discrete finite state machines and prescribed control laws, dEMG control used in this case study continuously drives a powered prosthesis joint based purely on the user’s neural control signals (i.e. motor commands) from the residual GAS and residual TA muscles. This device offered the amputee user the freedom to continuously adjust the behavior of prosthetic ankle (i.e. control both position and stiffness independently), which allows the amputee user to freely adapt their prosthesis behavior to accommodate versatile activities of daily life. We chose different postural control tasks during standing in this study, as the first step, to demonstrate the potential of dEMG control for standing postural control tasks that requires continuous coordination of residual muscle activation. Using preprogrammed autonomous control to accommodate versatile activities of daily life has been difficult because it requires the autonomous controller to seamlessly coordinate its behavior with a multitude of environments, contexts, and user intent.

One of the interesting observations in this study was that enabling neural control of a prosthetic ankle on the amputated side elicited improved motor coordination between the intact limb and amputated limb during postural control. The between-limb coordination was manifested by (1) synchronized CoP anterior-posterior excursion and (2) synchronized shank muscle activation. First, we observed a significant improvement in between-limb synchronization of CoP excursion during standing postural control when the TT amputee can actively use prosthetic ankle via neural control, compared to when he used passive device. Between-limb CoP synchronization has developed over recent years into a meaningful measure of postural control for populations with inter-limb deficits (i.e. stroke population) [33, 36]. When the participant wore a passive prosthesis, the missing ankle function led to lack of CoP excursion on the amputated side and therefore lack of bilateral CoP synchronization [37]. When the participant can actively move the ankle via the EMG control signals, not only the CoP excursion magnitude increased on the amputated side, but also it showed improved synchronization with the CoP excursion in the intact side. This CoP synchronization restores the possibility of normative CoP control strategies in standing typically observed in healthy individuals (i.e. CoP-CoM to CoM acceleration relationship [38]). The observation implies the importance in restoring ankle control and function for enhanced postural stability and the potential of dEMG control for active control of prosthetic ankle. Additionally, by demonstrating the ability for a transtibial amputee to volitionally adjust CoP excursion while improving standing postural control, this is the first study to show the potential for this biomechanical feature to indicate prosthetic ankle control capability. Second, the between-limb coordination was also observed in EMG activation pattern as shown in Figure 5. After learning the dEMG control of prosthetic ankle in standing postural control, nearly synchronized activation between intact and residual TA/GA was observed. One of the open questions is what neural mechanisms are responsible for the observed adaptation in residual muscle activations. The observation of synchronized activation in homologous muscles between limbs cause us to consider the potential for a common neural drive behind the activity for both muscles. It would be an interesting future direction to investigate the neuromuscular adaptation in lower limb amputees when the function of residual muscle activation is restored via dEMG control of prosthetic joints.

There is a lack of studies that have reported extended training and improvement in dEMG control in the lower limb with residual muscles. As a significant contribution, we developed a specific training paradigm guided by a physical therapist as well as acclimation strategy toward facilitating adaptation in residual antagonistic muscle activity. Instead of solely focusing on residual muscle activation for operation of prosthetic ankle (i.e. local joint level), our training also emphasized the full-body coordination and awareness in postural control while encouraging the participant in engaging prosthetic ankle via dEMG control during the task performance. Accordingly, over the course of training we witnessed various stages of learning from the amputee participant. During the initial training days (1 and 2) the amputee noted that he focused primarily on controlling the prosthetic ankle when completing the prescribed tasks. However, in the latter days of training (days 3-5) the participant frequently mentioned focusing on whole-body movement, using his prosthetic and intact limb symmetrically. Huang et al. observed improvement in dEMG control of a prosthetic ankle during walking when they provided visual feedback of the ankle-joint angle [27], demonstrating the relevance of this joint-level focus when learning. We extend the results from this study by demonstrating the ability for an amputee to potentially continue the learning process beyond this joint level focus, without the use of visual feedback. Since this learning occurred in the absence of supplementary artificial feedback, only under the guidance of verbal feedback from a physical therapist, this type of training shows promise toward real-world application of dEMG control of a powered ankle prosthesis. While the stages of learning observed here are discussed qualitatively, future investigations of amputee learning the dEMG control of a prosthetic device would benefit by analyzing the potential change in multi-joint muscle coordination via muscle synergy analysis [39, 40].

Our task-specific training, emphasizing both local joint control and full-body control, improved the participant’s postural control capability significantly. Before conducting this study, we did not know whether our recruited amputee participant could coordinate his residual muscle activation appropriately for prosthetic ankle control to assist postural stability. This was because in our previous studies [26], in which the amputee participant was also a test participant, he showed average performance compared with other transtibial amputees when asked to coordinate residual antagonistic muscles to balance a virtual inverted pendulum with human-like dynamics in a sitting position. In addition, it was unclear how the participant’s demographics, such as age (57y/o), BMI (~34), presence of vascular disease (including partial neuropathy at the intact foot), might affect his ability to improve control during training. Though these factors may have a significant negative effect on standing postural control [41–43] they are highly characteristic traits of the lower-limb amputee population [44, 45]. The before-training evaluation also showed limited muscle activation in residual TA (Figure 5) and comparable or even worse quiet standing test score (Table 2). However, when our training protocol was applied, we observed significant improvements in residual muscle control of the powered ankle prosthesis and postural control capability in both singular sessions and across training as a whole (Figure 4). Through this case study we have presented the first potential timeline for the improvement of dEMG control facilitated by PT-guided training over multiple days. Future study that wishes to accurately investigate the usefulness of dEMG control of lower-limb prostheses should consider the potential stage of learning of the individual amputees and the influence of improvement that can come with time and appropriate training.

The results from this study have several implications for the potential clinical benefit of dEMG control of a powered prosthetic ankle. During the follow-up evaluation of the load transfer task we observed the participant had limited range of motion with his passive prosthetic ankle, likely due to minimal change in angle of the stiff ankle joint. Hence, compensation with more trunk flexion was used, which is a known problem for back injuries during weightlifting. The participant was able to significantly change ankle angle using the dEMG control ankle allowing for an improved overall postural configuration (i.e. more vertical trunk angle) [46] in lifting, which could significantly prevent secondary injuries post amputation. In addition, this study has taken one of the first steps, via direct EMG control, toward addressing the normalization of other functional tasks (aside from locomotion) that are critical to daily life activities and amputee quality of life.

Our study included one amputee to investigate the feasibility of dEMG control of a powered ankle for enhanced postural control. Although exciting results were observed, this case study was insufficient to conclude the benefit of dEMG control of powered ankles. Future work should expand this current case study to include more participants to understand the applicability to the general amputee population. It would be interesting in future study for more measures of stability (including center of mass, joint torque symmetry, etc.) to further inform the effect of dEMG control of a powered ankle prosthesis. Though its effect is not specifically addressed in the context of this study, future study would benefit to evaluate the effect of prosthetic socket design on residual muscle activations during EMG control of lower-limb prostheses.

## Conclusion

This case study was the first attempt to demonstrate the feasibility and potential for direct EMG control of a powered prosthetic ankle, combined with PT-guided training, to enhance standing postural control across various contexts and tasks. The participant when using dEMG-controlled powered ankle yielded improved clinical balance score, reduced compensation from the intact joints, and improved between-limb coordination, compared to those when using his daily passive prosthesis. In addition, the case study developed a PT-guided training protocol for transtibial amputees, which is necessary for them in learning dEMG control of powered ankle to assist postural control and improve postural stability. This case study has developed the grounds for future design of versatile and agile powered lower-limb prostheses via direct, continuous EMG control via residual muscles, which may further improve the motor function of individuals with lower limb amputations and improve the ability for amputees to navigate standing postural control tasks that are a significant portion of daily-life activities.

## Declarations

### Ethics approval and consent to participate

The experimental protocol was approved by the Intuitional Review Board at the University of North Carolina – Chapel Hill. All participants provided written, informed consent.

### Consent for publication

The participant provided written, informed consent for publication.

### Availability of data and material

The datasets used and/or analyzed during the current study are available from the corresponding author on reasonable request.

### Authors’ contributions

SH, and HH contributed to the design of the experiment. SH, EB, and FH conducted the experiments. AF and SH processed the data. AF, SH, and HH wrote the manuscript. All authors contributed to the analysis and interpretation of the results and read and approved the final manuscript.

## Acknowledgements

The authors would like to acknowledge Abby Sweitzer and Loryn Boorstein for their help with data collection. We’d also like to thank all the undergraduate researchers and participants for their time and assistance with this research project.

